# Systematic integration of GATA transcription factors and epigenomes via IDEAS paints the regulatory landscape of mouse hematopoietic cells

**DOI:** 10.1101/730358

**Authors:** Ross C. Hardison, Yu Zhang, Cheryl A. Keller, Guanjue Xiang, Elisabeth Heuston, Lin An, Jens Lichtenberg, Belinda M. Giardine, David Bodine, Shaun Mahony, Qunhua Li, Feng Yue, Mitchell J. Weiss, Gerd Blobel, James Taylor, Jim Hughes, Doug Higgs, Berthold Gottgens

## Abstract

Members of the GATA family of transcription factors play key roles in the differentiation of specific cell lineages by regulating the expression of target genes. Three GATA factors play distinct roles in hematopoietic differentiation. In order to better understand how these GATA factors function to regulate genes throughout the genome, we are studying the epigenomic and transcriptional landscapes of hematopoietic cells in a model-driven, integrative fashion. We have formed the collaborative multi-lab VISION project to conduct **V**al**I**dated **S**ystematic **I**ntegrati**ON** of epigenomic data in mouse and human hematopoiesis. The epigenomic data included nuclease accessibility in chromatin, CTCF occupancy, and histone H3 modifications for twenty cell types covering hematopoietic stem cells, multilineage progenitor cells, and mature cells across the blood cell lineages of mouse. The analysis used the **I**ntegrative and **D**iscriminative **E**pigenome **A**nnotation **S**ystem (IDEAS), which learns all common combinations of features (epigenetic states) simultaneously in two dimensions - along chromosomes and across cell types. The result is a segmentation that effectively paints the regulatory landscape in readily interpretable views, revealing constitutively active or silent loci as well as the loci specifically induced or repressed in each stage and lineage. Nuclease accessible DNA segments in active chromatin states were designated candidate *cis*-regulatory elements in each cell type, providing one of the most comprehensive registries of candidate hematopoietic regulatory elements to date. Applications of VISION resources are illustrated for regulation of genes encoding GATA1, GATA2, GATA3, and Ikaros. VISION resources are freely available from our website http://usevision.org.

## Introduction

A person’s genetic profile can have a significant impact on complex traits such as disease susceptibility and response to specific treatments. Genome-wide association studies (GWAS) have mapped loci at which common genetic variation is associated with complex traits, but the mechanistic connection between genotype and phenotype is rarely understood. This is because most trait-associated genetic variants are not in the 1-2% of the genome that encodes mRNA, but rather in the much larger noncoding genome (1). While no DNA-based grammar has been developed yet to interpret these noncoding variants (2), the fact that they are highly enriched in chromatin with epigenetic features associated with gene regulatory elements offers new avenues to understanding their impact on phenotypes (3–5). Efforts to harvest GWAS results for potential medical application has led to the concept of precision medicine, in which a person’s genotype is used to improve lifestyle choices and develop therapeutic interventions specifically for that person (6). However, precision medicine requires more than genotypes and associations. Precision medicine needs a thorough understanding of the epigenome to interpret the large majority of trait-associated genetic variants that lie outside coding regions.

The problem we address is how to utilize the enormous amounts of emerging epigenetic data effectively both for basic research and precision medicine. Powered by advances in sequencing technologies, biochemical reagents, and bioinformatic analyses, many laboratories and large consortia, such as ENCODE (4), Roadmap Epigenome Project (7), GTEx (8), BluePrint (9, 10) and IHEC (11)) are determining transcriptome profiles and producing genome-wide views of the regulatory landscape (12). At this point, data acquisition may no longer the major barrier to understanding mechanisms of gene regulation during normal and pathological development. In fact, the volume of data produced is already overwhelming for most investigators. We seek to understand how epigenetic features regulate differentiation and how that regulation is altered in disease. Major challenges include integration of epigenetic data in terms that are accessible and understandable to a broad community of researchers, building validated quantitative models explaining how changes in epigenetic features affect the dynamics of gene expression across differentiation, and translation of the information effectively from mouse models to potential applications in human health.

Consider any genetic locus implicated in development, differentiation, behavior, or disease. Investigators may want to study the regulation of expression of gene(s) in that locus, e.g. to understand how genetic variants could affect its expression. This investigation could be greatly facilitated by abundant genome-wide datasets on multiple epigenetic features. Currently, to utilize such information, an investigator would examine epigenetic data around this locus in web-based genome browsers and databases. These resources are useful, but they do not cover all the relevant aspects of chromatin structure, dynamics, and expression. After finding the available data, the investigator will need to analyze the results to predict candidate *cis*-regulatory elements (cCREs), including enhancers, silencers, or insulators. While progress continues to be made in the predicting cCREs, issues of completeness (how sensitive are the cCREs for discovering true regulatory elements?) and specificity (how likely is it that the cCREs are true regulatory elements?) are actively debated. Developing more useful collections or registries of high quality cCREs is a major current need in functional genomics.

We have formed an interdisciplinary collaborative team to address these needs via **V**al**I**dated **S**ystematic **I**ntegrati**ON** of epigenomic data **(VISION)** to analyze and interpret molecular mechanisms regulating hematopoiesis in mouse and human. We are consolidating hundreds of epigenomic datasets and applying integrative approaches to generate robust candidate functional assignments to DNA segments. These assignments, coupled with gene target predictions and results of genome editing experiments, are the input to machine-learning approaches that generate quantitative models for how each candidate CRE contributes to the regulation of its target gene. Importantly, these models will be rigorously tested and validated by targeted genome editing in reference loci, and then applied genome-wide. Furthermore, we are developing resources to enable more accurate translation of regulatory insights between mouse and human. The results from our project will inform investigators about candidate CREs and their predicted roles in regulating their loci of interest, thus enabling them to design model-driven experiments to deepen their understanding of the investigated process.

In this concise review, we focus on our efforts to integrate the large amount of genome-wide information on epigenetic features and transcriptomes in a systematic manner to assign chromatin states across hematopoietic cells and predict cCREs. These resources are illustrated with respect to the GATA factors, both the genes encoding them and the binding patterns of the proteins in erythroid and lymphoid T-cells. A further examination of the *Ikzf1* gene encoding the Ikaros transcription factor illustrates the power of our integrative approaches to deduce data-driven hypotheses about differential regulation of gene expression in hematopoiesis.

### Compile and determine epigenetic features and transcript levels across hematopoietic differentiation

Over the past decade, the amount of information about gene expression levels and epigenetic regulatory landscapes in mammalian hematopoietic cells has increased exponentially, both through the work of individual laboratories (e.g. 13, 14-27) as well as the work of major consortia such as ENCODE and Blueprint. These data currently are provided in differing formats from diverse resources, with no common data processing or analysis, e.g. to find significant peaks of signals. Our first step in the VISION project was to compile the datasets, process the data in a consistent manner, and provide the data a manner enabling investigators to find all relevant information.

Building on resources independently developed in laboratories within the VISION project, we have established a distributed data network to enhance accessibility and develop a unified interface to the users. The **CODEX** resource, developed by the Gottgens group, maintains a compendium of next generation sequencing datasets pertaining to transcriptional programs of mouse and human blood development (28). The compendium currently contains over 1,700 publicly available datasets, all uniformly processed to facilitate comparisons across datasets. CODEX contains ChIP-seq, DNase-seq, and RNA-seq datasets, which are available as signal tracks, mapped sequence files, peak calls, and transcript levels for the RNA-seq. The CODEX website also provides a number of analysis tools including correlation analysis, sequence motif discovery, analysis of overrepresented gene sets, and comparisons between mouse and human. The **SBR-Blood** resource, developed by the Bodine lab, has compiled expression data, ChIP-seq, and Methyl-seq data for mouse and human hematopoietic cells (990 datasets), including normalizations across disparate datasets (29). Both of these resources feed into the **VISION** project, which provides raw and normalized datasets selected to cover specific groups of features in mouse and human hematopoiesis, segmentations by integrative modeling (see below), and catalogs of cCREs, among other resources, on the website http://usevision.org. This website includes a link to a genome browser with epigenetic and expression datasets during hematopoiesis as well as the **3D Genome Browser** developed by the Yue lab (30). In addition to the effort to compile and analyze existing data, new data are being generated both within the VISION project and in other laboratories that expand the coverage of epigenetic features across cell types and bring in datasets on new transcription factors or co-factors.

Our initial efforts were in mouse hematopoiesis because of the large number of epigenomic and transcriptomic datasets that were available in both primary maturing cells (exemplary references at the beginning of this section) and in the multilineage progenitors to blood cells (31). In addition, epigenomic data were included from selected cell lines that have been used extensively as models for multilineage myeloid cells (HPC7 cells, 32) and for GATA1-dependent erythroid maturation (G1E and G1E-ER4 cells, 33). The cell types investigated can be viewed in a simplified hierarchy (Figure 1A), which serves as a useful organizing structure despite the greater complexity discovered in single cell transcriptomes (34). For assignments to chromatin states, we focused on nuclease accessibility of chromatin, as determined by DNase-seq (35) or ATAC-seq (36), binding by the structural protein CTCF, and post-translational modifications of histone H3 N-terminal tails (37) associated with enhancers (H3K4me1), promoters (H3K4me3), active enhancers and promoters (H3K27ac), transcriptional elongation (H3K36me3), polycomb repression (H3K27me3), or heterochromatic repression (H3K9me3). For some cell types, all these features were determined (Figure 1B). Notably, the remaining cell types were missing data on multiple features. We stress that this problem with missing data is not unique to our work, but rather it is commonly seen in all large-scale analyses including Roadmap, ENCODE, and Blueprint. We suggest that the approach developed in VISION (see below) will be broadly useful in any setting with missing data. Estimates of transcription levels were available from RNA-seq in all the investigated cell types.

**Figure 1.**
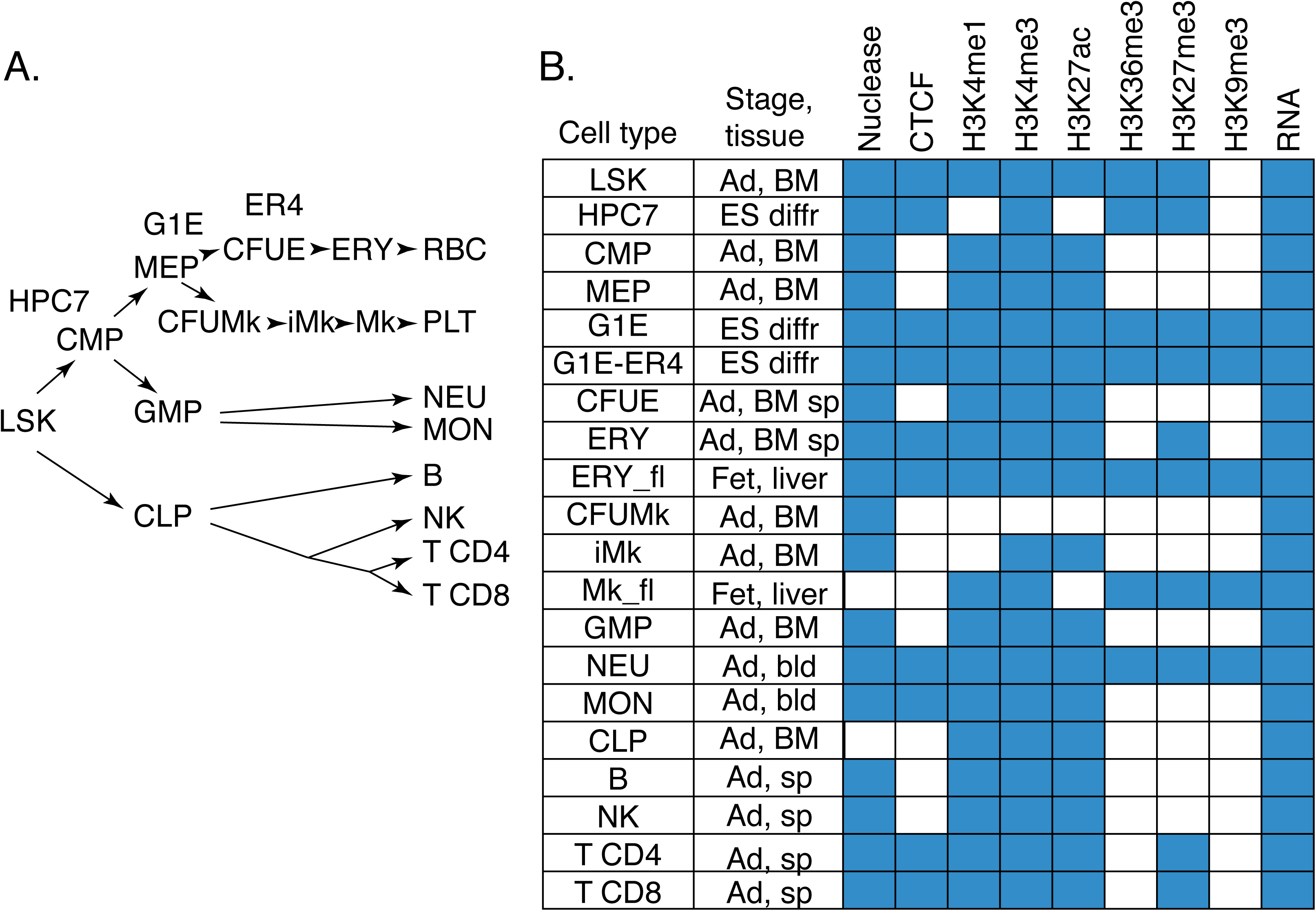
Mouse hematopoietic cell types and genome-wide datasets used in integrative analysis in the VISION project. **A.** The purified cell populations and cell lines are presented in a simplified hierarchical view. LSK= lineage minus, Sca1+, Kit+ cells, which include hematopoietic stem cells and early multilineage progenitor cells; CMP=common myeloid progenitor cells; MEP=megakaryocytic erythroid progenitor cells; CFUE=colony-forming units erythroid; ERY=erythroblasts, RBC=red blood cells; CFUMk= colony-forming units megakaryocytic; iMk=immature megakaryocytic cells; Mk= megakaryocytes; PLT=platelets; GMP=granulocyte monocyte progenitor cells; NEU=neutrophils; MON=monocytes; CLP=common lymphoid progenitor cells; B=lymphoid B cells; NK=natural killer cells; T CD4=CD4+ T-cells; T CD8=CD8+ T-cells; HPC7=cell line that serves as a model of a multipotent myeloid progenitor cell; G1E and ER4=cell lines that serve as a model for GATA1-dependent erythroid maturation. The G1E cell line was derived from *Gata1-* ES cells induced toward erythroid differentiation; they are blocked from further differentiation at an early stage after erythroid commitment. ER4 refers to the sub-line G1E-ER4, which expresses an inactive hybrid of GATA1 fused to the hormone binding domain of the estrogen receptor; these cells show rapid maturation toward late stage erythroblast-like cells upon estradiol treatment. **B.** Epigenomic and transcriptomic datasets available for each cell type are indicated by a blue-filled box in the matrix. Ad=adult; Fet=fetal; ES=embryonic stem cells; BM=bone marrow; sp=spleen; bld=blood; diffr=differentiated.

### Systematically learn and assign epigenetic states across cell types

The large number of interrelated epigenetic datasets described above present immense opportunities for understanding differential gene regulation, if these data can be integrated into robust annotation of likely functional DNA. A key challenge is to build quantitative models explaining how the dynamics of epigenomes across many cell types lead to gene expression changes and phenotypic diversity (4, 38). A current approach for describing epigenetic landscapes is genome segmentation (39, 40), which assigns states to genomic segments exhibiting unique patterns of chromatin marks. Existing genome segmentation tools (39–42) were developed primarily for segmenting the epigenomes of single cell types. Although genomes from different cell types may be concatenated together, such an approach ignores the position-specific epigenetic events conserved across related cell types. We have used the IDEAS method (**I**ntegrative and **D**iscriminative **E**pigenome **A**nnotation **S**ystem) for two-dimensional segmentation along chromosomes and across cell types because of its improved accuracy and consistency in assigning epigenetic states (43, 44). This method uses a Bayesian model to approximate quantitative data distributions without signal binarization. It also utilizes Bayesian techniques to automatically determine the best model sizes, including the number of states.

Importantly, the statistical framework for IDEAS allows it to assign likely epigenetic states to cell types based on the data distributions for signals across cell types. Thus, even if particular features have not been determined in a cell type, the system can still assign a likely state based on the known signals in other, locally related cell types. This model-based inference of states has better performance than current data imputation procedures (45). As noted above (Figure 1B), many of the cell types of interest did not have data on all features, but we were able to utilized this ability of IDEAS to produce segmentations despite missing data to generate informative segmentations across all the cell types examined.

The IDEAS segmentation method is analogous to integration by mixing. One can consider the signal track for each epigenetic feature as a signal with a distinctive color, as illustrated in Figure 2 with deep red for DNase-seq, purple for CTCF, etc. An intuitive way to integrate the eight tracks of information is simply by mixing, e.g., by merging all the colored tracks into one. That approach does bring out some aspects of the combined features, such as CTCF and nuclease accessibility 5’ to *Zfpm1* and a mix of K4 monomethylation and K36 trimethylation of H3 through the body of the gene. However, the mixing can also blend too many colors together to distinguish clear states, such as around the transcription start site (TSS) and the region around the 3’ end of *Zfpm1*. The systematic integration by segmentation can be thought of as a principled, objective way to find well-defined, discrete combinations of features that occur frequently throughout the epigenomes examined (the epigenetic states). Each genomic segment is then assigned to the one state that best matches the known (or inferred) epigenetic signals in each cell type. Thus, the IDEAS track below “merge tracks” gives a principled resolution of the mixtures of the epigenetic features.

**Figure 2.**
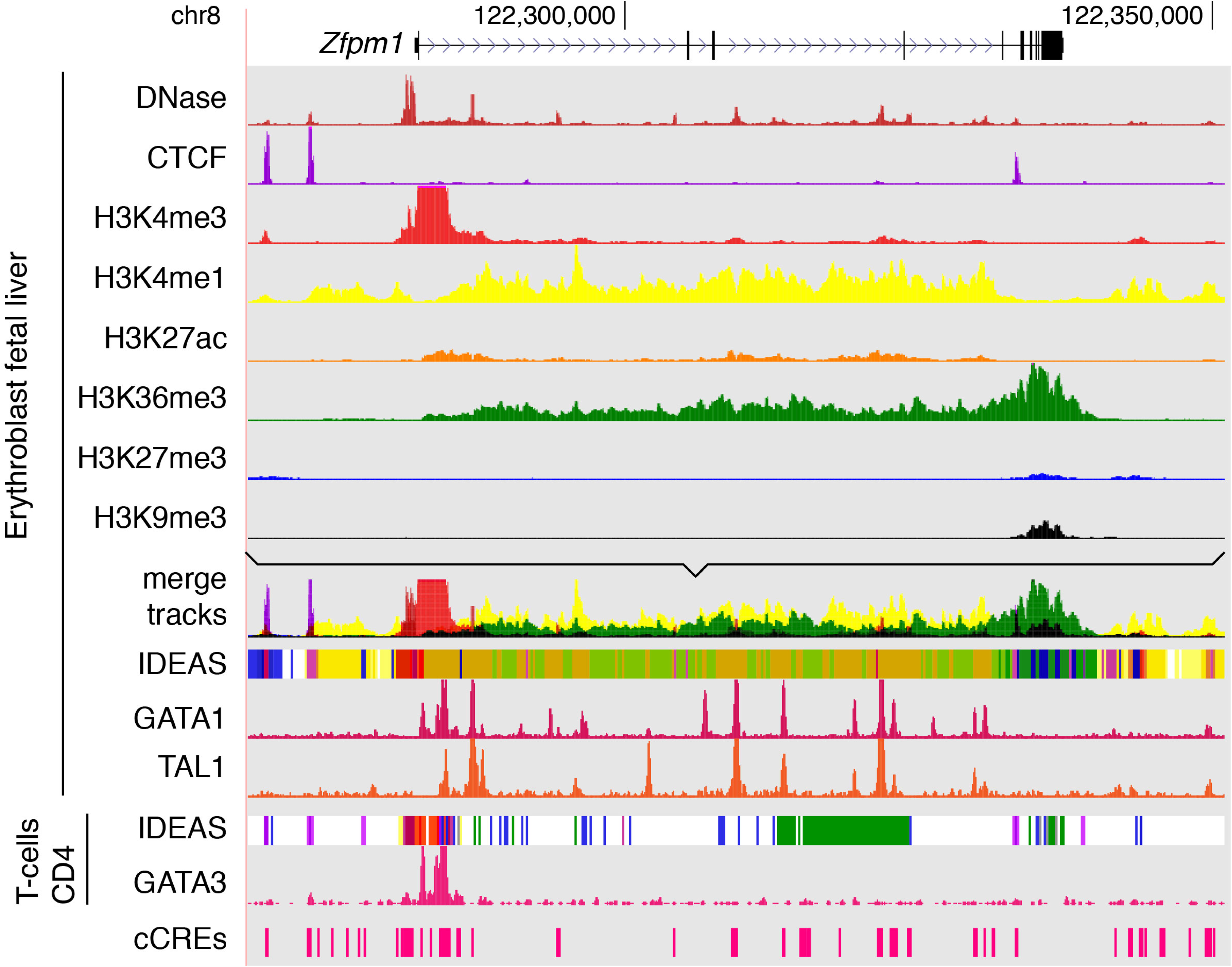
Merging and systematic integration of epigenetic data tracks. For an 83 kb region around the *Zfpm1* gene (GRCm38/mm10 assembly; chr8:122,268,001-122,351,000), the tracks in the VISION genome browser (http://usevision.org) show the gene model; signal tracks for eight epigenetic features, each with a distinctive color, in fetal liver erythroblasts; a merge of the eight tracks (using the Track Collections tool from the UCSC Genome Browser); the IDEAS segmentation of the epigenetic signals after normalization; ChIP-seq signal tracks for GATA1 and TAL1 in fetal liver erythroblasts; the IDEAS track and GATA3 ChIP-seq for CD4+ T-cells; and the VISION cCREs.

For the eight epigenetic features across 20 mouse hematopoietic cell types, IDEAS generated a 27-state model (Figure 3). Each state was defined by a quantitative profile of signal strengths from the features, illustrated by the heat map. Each of the eight features was assigned a color, and these were used in turn to establish automatically a color for each state based on the contribution of each feature to that state. Thus, the several promoter-like states were colored in various shades of red, the enhancer-like states were given yellow to orange colors, CTCF-containing states had purple colors, transcribed states were colored in states of green, states associated with polycomb-repression were blue, the heterochromatic state was gray. The most frequently occurring state (state 0) was a quiescent state, with very low signal for each of the eight features. Many of these combinations of features have been described previously, and the segmentation provided a systematic means to identify those combinations as states that are assigned consistently across cell types. The IDEAS states also gave a discrete set of several promoter-associated or enhancer-associated states, which can be further examined experimentally for functional roles.

**Figure 3.**
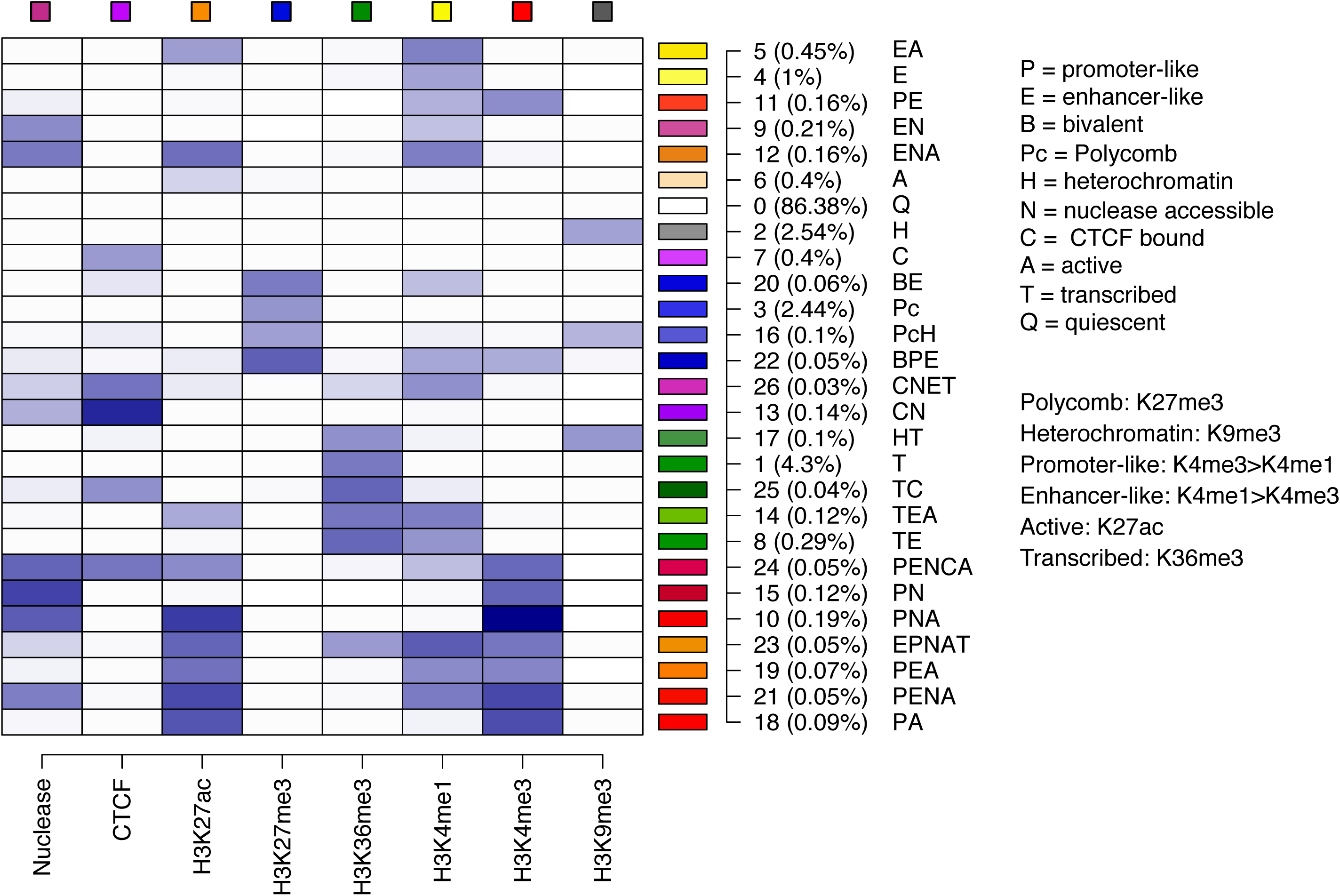
Epigenetic states learned by IDEAS modeling. The 27 epigenetic states (rows in the matrix) learned by the IDEAS system from the data on eight epigenetic features (columns in the matrix) are shown in a white-to-blue heatmap, with darker blue indicating stronger contribution of a feature to a state. To the right of each row is shown the color and number for each state, the average percentage coverage of the genomes in the 20 cell types by that state, and a label for each state consisting of the one or two letter abbreviations for the function-associated attributes listed in the key at the right.

The epigenomic data were determined in many different laboratories at different times, with systematic differences in protocols, sequencing depths, and other factors that could impact the integrative analysis. Thus, consistency in data processing and appropriate normalization of the data were also key components of the IDEAS segmentation pipeline. More complete descriptions of the approaches developed and utilized are given elsewhere (46, 47). The impact of normalization is illustrated in Figure 2. The upper tracks of individual features show the raw numbers of reads mapped to DNA intervals for one of two replicates, all set to the same scale. However, after normalizing to adjust for differences in sequencing depth and in the signal-to-noise ratio, the H3K27me3 signal was boosted such that it drove the assignment of some DNA segments at the left end of the diagram to the polycomb repressed (blue) state.

The accuracy and effectiveness of the IDEAS segmentation can be evaluated by comparison to orthogonal data that provides an alternative view of the functions implied by the segmentation. Specifically, we examined the binding patterns of GATA transcription factors known to regulate gene expression in different hematopoietic lineages. The transcription factors GATA1, GATA2, and GATA3 have been strongly associated with enhancers and transcriptional switches in erythroid, myeloid progenitor cells, and lymphoid cells, respectively (48). Thus, one may expect the enhancer-like states in the *Zfpm1* gene in erythroblasts to be bound by GATA1. Indeed, the ChIP-seq pattern for GATA1 coincided well with those states (orange in the IDEAS track, Figure 2). Moreover, these enhancer-like states were co-bound by the transcription factor TAL1 in erythroblasts (Figure 2). This co-binding by GATA1 and TAL1 has been strongly associated with gene activation (17, 18, 49), and the ChIP-seq patterns for these transcription factors lend strong support to the epigenetic state assignments. Many of these predicted enhancers of *Zfpm1* were shown to increase expression from a reporter gene in transfected cells, giving further credence to the segmentation results (25, 50).

By conducting the segmentation jointly across cell types as well as along chromosomes (the two-dimensional segmentation), the IDEAS method also brings out differences between cell types. The segmentation of *Zfpm1* in CD4+ T-cells differed greatly from that in erythroblasts (Figure 2). Many of the DNA segments in the gene body that were enhancer-like in erythroblasts were either in the quiescent (white) or transcribed (green) states in CD4+ T-cells. The region around the TSS was assigned to promoter-like (red states). Notably, this same TSS region is bound by GATA3 in CD4+ T-cells, as indicated by the ChIP-seq signal (obtained from CODEX for data from 51). Thus, expression of *Zfpm1* appears to be regulated at the TSS in CD4+ T-cells, whereas multiple internal enhancers are utilized in erythroid cells.

### Define a large set of cCREs in mouse hematopoietic cells

The integrative segmentation from IDEAS allowed us to take a straightforward approach to predicting candidate *cis*-regulatory elements or cCREs (47). The nuclease-accessible DNA intervals (hypersensitive sites or HSs) in each cell type were determined by peak-calling method on the DNase-seq and ATAC-seq data. We then gathered all HSs from all cell types (requiring replication within a cell type if available) and merged the overlapping ones. This set of HSs was then filtered to remove any that were only in the quiescent state (0) in all cell types. The remaining set contained all HSs that were in an IDEAS state indicative of dynamic histone modifications or CTCF binding in at least one of the cell types examined. This simple two-step method for predicting cCREs relies on the sophistication of IDEAS for assigning DNA intervals to one of the commonly occurring combinations of epigenetic features. It does not rely on any particular combination of histone modifications to predict cCREs, and it should be robust to changes in epigenetic landscape that result from switches in regulatory and expression patterns between cell types.

The initial registry of cCREs was comprised of 205,019 DNA intervals in eighteen hematopoietic cell types in mouse (no nuclease sensitivity data were available for two of the twenty cell types, Mk from fetal liver and CLP). The absence of full knowledge of functional elements and neutral elements across genome precludes a rigorous determination of the sensitivity and specificity of this initial cCRE registry. However, this collection does look promising in several respects. The registry captured virtually all the known erythroid cCREs, and it included a large majority (two-thirds) of DNA segments bound by the transcription co-activator EP300 in murine erythroleukemia cells, CH12 cells (a model for B cells), and fetal liver (47). Thus, the recall appears to be reasonable. Further experimental tests should provide insight into the precision or specificity of the cCRE predictions.

The cCREs in and around the *Zfpm1* gene are shown on the bottom line of Figure 2. They include the candidate enhancer-like regions discussed above, as expected. Many other cCREs are also present, which raises questions such as (1) in what cell types does a particular cCRE appear to be active? (2) What transcription factors may be bound to a cCRE? (3) How likely is it that a cCRE is regulating a gene of interest? Continuing work in the VISION project strives to address such questions. Question 1 is addressed by the state assignments for each cCRE in each cell type, which can be downloaded or browsed at the project website. Question 2 is being addressed by utilizing the resources in CODEX to annotate cCREs with binding data from ChIP-seq. Question 3 is being addressed by developing quantitative models to “explain” gene expression data in terms of IDEAS state assignments across cell types. Future work should bring in chromatin interaction data and additional machine learning approaches.

### Illustrate the VISION resources at loci encoding GATA factors and Ikaros

As discussed above, the GATA family of transcription factors is well-known for regulation of gene expression in specific cell types and lineages. Examination of the genes encoding these factors and another key regulator of gene expression in hematopoietic cells, Ikaros (IKZF1), illustrates the types of insights that investigators can glean from the integrative analyses in the VISION project. Levels of expression of the genes were estimated from the RNA-seq data from recent publications (27, 31) that were compiled at the VISION web site (Figure 4A). Consistent with the lineage specificity previously reported, expression of *Gata1* was most prevalent in erythroid cells and the multilineage progenitor cell populations CMP and MEP, with more modest expression in megakaryocytic cells. The *Gata2* gene was expressed more highly in the multilineage progenitor cell populations with some persistence into megakaryocytic cells. High levels of *Gata3* expression were found primarily in a subset of the lymphoid cells, *viz*. NK, CD4+ and CD8+ T-cells. In contrast, expression of *Ikzf1* was expressed at higher levels and in a broader pattern, with expression in most hematopoietic cell types albeit lower in maturing erythroid cells.

**Figure 4.**
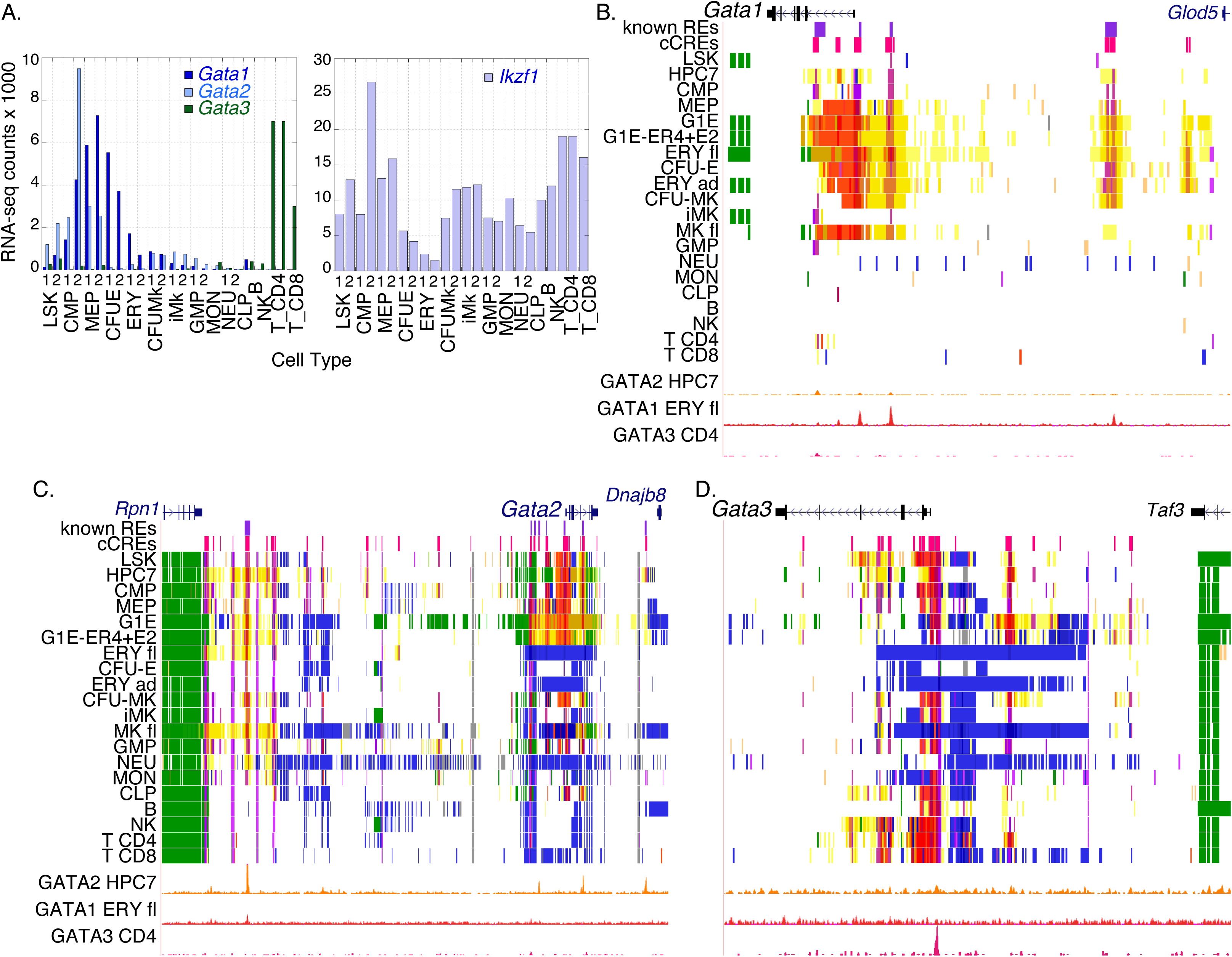
Expression levels and epigenetic landscapes for genes encoding hematopoietic GATA factors. **A.** Estimated expression levels for the *Gata1*, *Gata2*, and *Gata3* genes (*left* graph) and the *Ikzf1* gene (*right* graph). RNA-seq determinations were in duplicates in most cell types, indicated by the digits 1 and 2. **B. – D.** Epigenetic landscapes are shown as genome browser views of gene models, known erythroid CREs (purple rectangles), predicted cCREs (bright red rectangles), IDEAS segmentation tracks in 20 cell types, and binding profiles for GATA2 in HPC7 cells, GATA1 in fetal liver erythroblasts, and GATA2 in CD4+ T-cells. **B.** A 50 kb region around the *Gata1* gene (GRCm38/mm10 assembly; chrX:7,955,001-8,005,000). **C.** A 130 kb region around the *Gata2* gene (chr6:88,095,001-88,225,000). **D.** A 70 kb region around the *Gata3* gene (chr2:9,850,001-9,920,000).

The epigenetic landscapes summarized as states from the IDEAS model showed patterns that fit with the cell type specificity of expression, and they revealed potential regulatory elements (cCREs) involved in cell type-specific control of expression. The IDEAS tracks around *Gata1* showed active epigenetic states in MEP, erythroid, and megakaryocytic cells (Figure 4B), which also express this gene. Furthermore, this locus has six cCREs, four of which have been shown to be enhancers or promoters regulating *Gata1* expression in erythroid cells (52, 53). Both known CREs and novel cCREs were bound by GATA1 in erythroid cells, albeit at varying levels, but none were bound by GATA2 in the myeloid progenitor cell model HPC7 cells or by GATA3 in CD4+ T-cells. In cell types not expressing *Gata1*, the locus was largely in the quiescent state, indicating that histone H3 in the chromatin of these cell types was undergoing little to no dynamic modifications.

Several regulatory elements have been mapped in the *Gata2* locus, both proximal and internal to the gene as well as distal, close to the *Rpn1* gene (50, 54, 55). These regulatory elements were in active epigenetic states in the expressing cell types, and the distal CRE was in an active state in a broader range of cell types (Figure 4C). Several of the CREs were bound by GATA2 in HPC7 cells (as well as the previously reported binding in G1E cells, not shown), but little to no binding was observed for GATA1 or GATA3. In contrast to the quiescent state observed for non-expressing cells for the *Gata1* locus, the *Gata2* locus was in a polycomb repressed state (H3K27me3) in many of the non-expressing cells. These distinct mechanisms inferred for repression (quiescent vs. polycomb) were deduced simply by examining the IDEAS tracks, and they illustrate insights that follow easily from integrative analysis and modeling.

The DNA interval around the TSS of the *Gata3* gene was in an active promoter-like epigenetic state and was bound by GATA3 in lymphoid cells, consistent with the expression pattern (Figure 4D). However, several additional DNA segments internal to the gene and upstream (between *Gata3* and *Taf3*) were in active states and were inferred to be cCREs. Thus, the regulation of *Gata3* may involve multiple CREs. As with the *Gata2* locus, the *Gata3* locus tended to be in a polycomb repressed state in many non-expressing cell types. Surprisingly, the cCREs around *Gata3* were in active epigenetic states in multilineage progenitor cells such as LSK, despite the very low levels of expression. This apparently precocious activation of the epigenetic landscape may serve as a type of lineage priming, or it could reflect some lineage commitment in this cell population.

The epigenetic states and transcription factor binding around the more widely expressed *Ikzf1* revealed patterns indicative of lineage-specific regulatory mechanisms (Figure 5). In addition to the transcribed states internal to the gene in almost all cell types, multiple cCREs in active states were observed around the TSS, upstream to the gene, in the third and seventh introns, and downstream. Strikingly, the pattern of binding of GATA factors was lineage-specific, with GATA2 binding at an upstream cCRE and in intron 3 in multilineage progenitors, GATA1 binding at a different set of cCREs upstream and in intron 3 in erythroid cells, and GATA3 binding in still a different pattern in CD4+ T-cells. The cCREs tended to be in active enhancer-like or promoter-like states in the cell types for which binding by GATA factors was also observed. These distinct GATA binding patterns, coupled with active epigenetic states from IDEAS, indicate substantial lineage specificity in the cCREs and transcription factors utilized to achieve an appropriate level of expression of *Ikzf1* in the various hematopoietic cell types.

**Figure 5.**
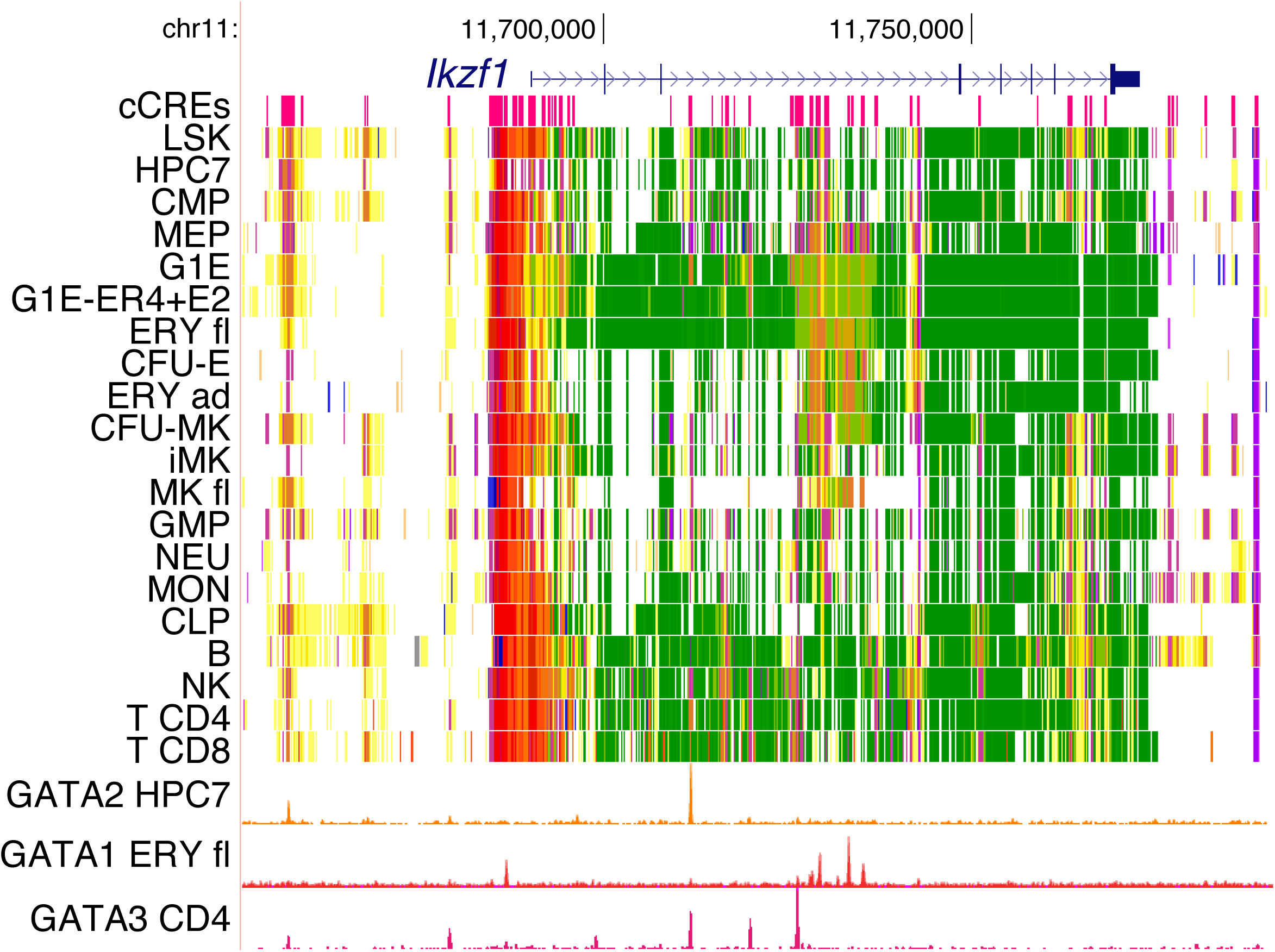
Epigenetic landscapes for the genes encoding Ikaros. For an 140 kb region around the *Ikzf1* gene (GRCm38/mm10 assembly; chr11:11,651,001-11,791,000), the tracks of cCREs, IDEAS states, and GATA factor occupancy are shown as in Figure 4.

### Future Perspectives

A major goal of the VISION project is to provide integrated views of epigenomic landscapes and transcriptomes from mammalian hematopoietic cells that will inform gene regulatory models to advance our understanding of global gene regulation. Importantly, these integrative views should enable other investigators to formulate data-based, testable hypotheses to advance their specific research interests. This review has focused on our recent work with mouse hematopoietic cells, organizing and analyzing about 150 tracks of epigenomic data from 20 cell types to produce a segmentation into well-defined epigenetic states using the IDEAS method. The consistent and distinctive colors associated with each state present the segmentation results as a type of painting, with one multi-colored panel for each cell type. Thus enhancer-like and promoter-like elements can be easily seen in a genome browser, as well as changes in the states among cell types. The epigenetic state assignments were used to annotate nuclease HSs and produce an initial registry of cCREs in mouse hematopoietic cells. This set of slightly over 200,000 cCREs serves as a large set of candidate regulatory elements that can be used in many ways for further research. Likely regulatory elements are now readily available for any gene, along with information about the epigenetic state of the chromatin covering that gene in the 20 hematopoietic cell types. The registry of cCREs can be examined for overlaps with lists of peaks of transcription factor binding (from ChIP-seq) for further inferences about potential functions of the cCREs.

Building from these initial resources, we have now compiled a large number of epigenomic datasets on human blood cells from the IHEC Blueprint Consortium (10) and many individual laboratories, including recent data on multilineage progenitor cells (56). These data are being integrated via IDEAS segmentation, and an initial registry of cCREs is being built using the approaches described here for mouse hematopoietic cells. In addition to the purposes already discussed, these resources will be particularly valuable for improving the interpretation of human genetic variants associated with various blood cell traits and diseases. Large scale GWAS have revealed many variants associated with traits of interest in hematology, and we now expect that many of the causative variants are acting through impacts on gene regulation (57). Having a set of high quality cCRE predictions decreases the search space for likely functional variants. Thus, the cCRE predictions may enable more precise, higher resolution studies of the potential impacts of the trait-associated, non-coding genetic variants.

A continuing challenge for utilizing cCRE predictions is the ambiguity in inferring a target gene. Regulatory elements can be far away from their target gene, and it is not uncommon for a CRE to be separated from its target gene by multiple non-target genes. Substantial efforts within the VISION project and elsewhere are tackling this enduring challenge. Measurements of chromatin interaction frequencies in an all-against-all mode such as Hi-C (58) or using capture strategies to focus on particular regions or interactions (23, 59) should provide important information to leverage with respect to target gene assignments. We are currently utilizing high resolution Hi-C data (26) and capture-C data (23, 60) from erythroid cells for multiple studies including improvement of target gene assignments. Furthermore, we are using multivariate regressions (47) and machine learning approaches (61) to estimate the impact of individual cCREs on potential target genes. While such studies can be limited with respect to statistical power, we are developing approaches that make effective use of the available data. In fact, one of the several important outcomes from our VISION project to help motivate new methods development to provide robust results to drive further research.

All resources from the VISION project are publicly available via our website http://usevision.org. We hope that this review will encourage use of these resources.

## Acknowledgements

The VISION project is supported by a grant from NIH/NIDDK R24DK106766 (multi-PI). Additional support of this work is from NIH/GM R01GM121613 (to Y.Z. and S.M.), NIH/NIDDK R01DK054937 (to G.B.), and NIH/NCI R01CA178393 (to R.H.).

